# Homotypic and heterotypic *trans-assembly* of human Rab-family small GTPases in reconstituted membrane tethering

**DOI:** 10.1101/544379

**Authors:** Kazuya Segawa, Naoki Tamura, Joji Mima

## Abstract

Membrane tethering is a highly regulated event that occurring during the initial physical contact between membrane-bounded transport carriers and their target subcellular membrane compartments, thereby ensuring the spatiotemporal specificity of intracellular membrane trafficking. Although Rab-family small GTPases and specific Rab-interacting effectors, such as coiled-coil tethering proteins and multisubunit tethering complexes, are known to be involved in membrane tethering, how these protein components directly act upon the tethering event remains enigmatic. Here, using a chemically defined reconstitution system, we investigated the molecular basis of membrane tethering by comprehensively and quantitatively evaluating the intrinsic capacities of 10 representative human Rab-family proteins (Rab1a, -3a, -4a, -5a, -6a, -7a, -9a, -11a, -27a, and -33b) to physically tether two distinct membranes via homotypic and heterotypic Rab-Rab assembly. All of the Rabs tested, except Rab27a, specifically caused homotypic membrane tethering at physiologically relevant Rab densities on membrane surfaces (e.g., Rab-to-lipid molar ratios of 1:100-1:3000). Notably, endosomal Rab5a retained its intrinsic potency to drive efficient homotypic tethering even at concentrations below the Rab-to-lipid ratio of 1:3000. Comprehensive reconstitution experiments further uncovered that heterotypic combinations of human Rab-family isoforms, including Rab1a/6a, Rab1a/9a, and Rab1a/33b, can directly and selectively mediate membrane tethering. Rab1a and Rab9a in particular synergistically triggered very rapid and efficient membrane tethering reactions through their heterotypic *trans*-assembly on two opposing membranes. In conclusion, our findings establish that, in the physiological context, homotypic and heterotypic *trans*-assemblies of Rab-family small GTPases can provide the essential molecular machinery necessary to drive membrane tethering in eukaryotic endomembrane systems.

## Introduction

In eukaryotic endomembrane systems, biological molecules such as proteins and lipids are selectively sorted into membrane-bounded transport carriers (e.g., secretory and endocytic transport vesicles) as cargo molecules and specifically delivered to their appropriate subcellular locations (e.g., subcellular organelles, the plasma membrane, or the extracellular space) [1]. This process, termed membrane trafficking, is fundamental and essential in all eukaryotic cells and is achieved in a spatiotemporally regulated manner in the following sequence: membrane-bounded transport carriers containing the appropriate cargoes are (i) formed from donor membrane compartments, (ii) conveyed along actin and microtubule cytoskeletons (i.e., cytoskeletal transport), (iii) reversibly tethered to acceptor membrane compartments (i.e., membrane tethering), (iv) stably and irreversibly docked to acceptor compartments (i.e., membrane docking), and finally (v) fused with acceptor compartments to deliver their cargoes (i.e., membrane fusion) [1]. During these events in intracellular membrane trafficking, membrane tethering provides the initial physical contact between membrane-bounded transport carriers and their target subcellular membrane compartments, thus primarily contributing to the compartmental specificity of membrane trafficking [2–8]. A large body of earlier genetic and biochemical studies have reported that Rab (Ras [rat sarcoma] related in brain) family small GTPases and specific sets of Rab-interacting proteins (i.e., Rab effectors), such as the coiled-coil tethering proteins and multisubunit tethering complexes, participate in the membrane tethering process [2–11] that occurs prior to the membrane docking and fusion events mediated by SNARE (soluble NSF [N-ethylmaleimide-sensitive factor] attachment protein receptor) family proteins and SNARE-interacting chaperones [12, 13], which are additional critical steps to confer fidelity to the membrane trafficking process [12–17]. Although Rab-family small GTPases and specific Rab-interacting effector proteins have generally been recognized as the conserved protein families responsible for membrane tethering [18, 19], the mechanisms by which these key components directly and physically act upon the tethering process remain elusive [8, 20]. Using a chemically defined reconstitution system with purified protein components and synthetic liposomal membranes, recent biochemical studies have uncovered a handful of protein machineries that directly trigger membrane tethering, including the human Golgi coiled-coil protein GMAP-210 (Golgi microtubule-associated protein 210) [21], the human endosomal coiled-coil protein EEA1 (early endosome antigen 1) [22], the yeast heterohexameric HOPS (homotypic vacuole fusion and vacuole protein sorting) tethering complex [23–25], and Rab-family small GTPases in yeast [26] and humans [8, 27, 28]. Although the experimental data obtained from the chemically defined systems provide novel mechanistic insights into the molecular functions of these putative membrane tethers [8, 21–28], an important caveat is that the reconstituted tethering reactions observed *in vitro* can adequately recapitulate membrane tethering events that occur between the membrane-bounded carriers and subcellular membrane compartments *in vivo*, particularly in terms of specificity and efficiency [8]. In this study, to further explore the physiological significance of reconstituted Rab-mediated membrane tethering reactions [8, 27, 28], we comprehensively and quantitatively evaluated the intrinsic tethering potency of representative human Rab-family small GTPases to directly and physically link the two distinct lipid bilayers together via *trans*-assembly of membrane-anchored Rab proteins in homotypic and heterotypic arrangements.

## Results and discussion

Out of over 60 protein isoforms of Rab-family small GTPases in human cells, which constitute the largest branch of the Ras superfamily [29], we selected 10 representative human Rab-family proteins (Rab1a, Rab3a, Rab4a, Rab5a, Rab6a, Rab7a, Rab9a, Rab11a, Rab27a, and Rab33b; Figure 1, S1), to assess intrinsic membrane tethering capacities in this reconstitution study. All 10 of these human Rabs have been well defined with respect to their intracellular locations, which cover most of the major subcellular compartments in eukaryotic cells (Figure 1A), and their involvement in the secretory and endocytic trafficking pathways, as summarized in prior review articles [18, 19] and the UniProt protein knowledgebase (UniProtKB) [30]. They also all share a typical structural feature of the Rab small GTPase family, and are thus known to be small monomeric proteins of around 25 kDa that consist of an N-terminal non-conserved flexible segment, a conserved globular Ras-superfamily GTPase domain (G-domain) in the middle, a C-terminal flexible hypervariable region (HVR), and one or two geranylgeranyl lipid anchors, which are post-translationally conjugated to the C-terminus of the HVR (Figure 1A, S1) [8, 31, 32]. The current chemically defined reconstitution systems for human Rab GTPase-mediated membrane tethering mimics the membrane-bound state of native Rab proteins bearing the C-terminal geranylgeranyl lipid anchors inserted into the cytosolic surfaces of subcellular membranes [8, 18, 19]. This was achieved using recombinant proteins of the selected Rab-family proteins that were purified in their full-length forms with an artificially modified polyhistidine tag (His12) at the C-terminus (Figure 1A, B), which exhibits high affinity binding to a DOGS-NTA (1,2-dioleoyl-sn-glycero-3-{[N-(5-amino-1-carboxypentyl) iminodiacetic acid]-succinyl}) lipid that was used for the preparation of synthetic liposomal membranes (Figure 2–8) [8, 26–28]. In addition to the Rab-His12 proteins, we also prepared polyhistidine-tagged full-length forms of human HRas and Arf1 (Figure 1A, B) for use as control proteins of the “non-Rab” Ras superfamily small GTPases [29]. Because the native Arf1 is post-translationally modified by a myristoyl lipid group at the N-terminus [33], Arf1 was purified as the N-terminal His12-tagged form (His12-Arf1), while HRas contains farnesyl and palmitoyl lipid anchors at the C-terminus *in vivo* [34], and thus was purified as the C-terminal His12-tagged form (HRas-His12), which was the same form as the Rab-family proteins (Figure 1A). Before testing reconstituted membrane tethering of the synthetic liposomes in the presence of the prepared small GTPase proteins, purified Rab-His12, HRas-His12, and His12-Arf1 proteins were characterized by GTP-hydrolysis activity assays (Figure 2A) and liposome co-sedimentation assays (Figure 2B, C). All of the tested His12-tagged Rab-family and other small GTPase proteins retained their intrinsic hydrolytic activities to specifically convert GTP to GDP and a free phosphate (Figure 2A), which were comparable between the His12-tagged proteins and also with those of untagged Rab1a and Rab5a proteins lacking a polyhistidine tag (Figure 1, Figure 2A). Liposome co-sedimentation assays were employed with DOGS-NTA-containing liposomes (1000 nm in diameter) and purified Rab-His12, HRas-His12, His12-Arf1, and untagged Rab proteins (Figure 2B, C), indicating that the His12-tagged small GTPase proteins were efficiently and stably associated with the liposomal membranes, whereas no membrane association was observed with the untagged Rab1a and Rab5a (Figure 2C). These biochemical analyses confirmed that the purified Rab-family, HRas, and Arf1 proteins in the current preparations were correctly folded and functional proteins with native-like active G-domains, and that their artificial C-terminal or N-terminal His12 tags were able to support stable membrane attachment without affecting the functions of the G-domains.

**Figure 1.**
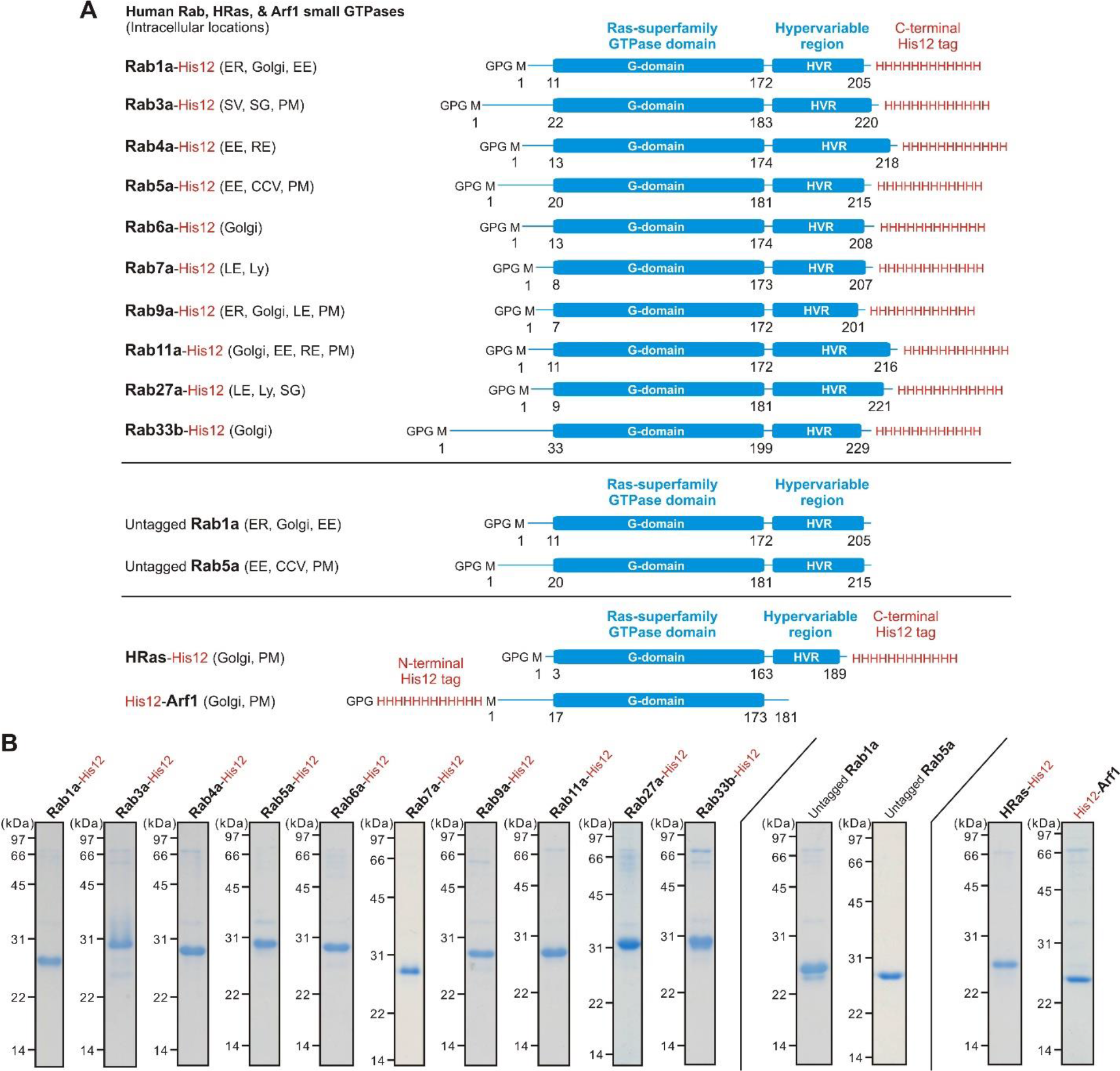
Human Rab-family small GTPases used in the current reconstituted proteoliposome systems. (**A**) Schematic representation of the C-terminal His12-tagged forms of human Rab-family small GTPases (Rab1a-His12, Rab3a-His12, Rab4a-His12, Rab5a-His12, Rab6a-His12, Rab7a-His12, Rab9a-His12, Rab11a-His12, Rab27a-His12, and Rab33b-His12), the tagless forms of human Rabs (untagged Rab1a and untagged Rab5a), and the C- or N-terminal His12-tagged forms of human non-Rab Ras-superfamily GTPases (HRas-His12 and His12-Arf1), showing their amino acid residues, domains (Ras-superfamily GTPase domains and hypervariable regions), and intracellular locations. The intracellular locations indicated include endoplasmic reticulum (ER), Golgi apparatus (Golgi), early endosome (EE), secretory vesicle (SV), secretory granule (SG), plasma membrane (PM), recycling endosome (RE), clathrin-coated vesicle (CCV), late endosome (LE), and lysosome (Ly). All of the recombinant small GTPases have only three extra N-terminal residues (Gly-Pro-Gly) after removal of the N-terminal GST-His6 affinity tags by proteolytic cleavage. (**B**) Coomassie blue-stained gels of purified Rab-His12, untagged Rab, HRas-His12, and His12-Arf1 proteins used in this reconstitution study.

**Figure 2.**
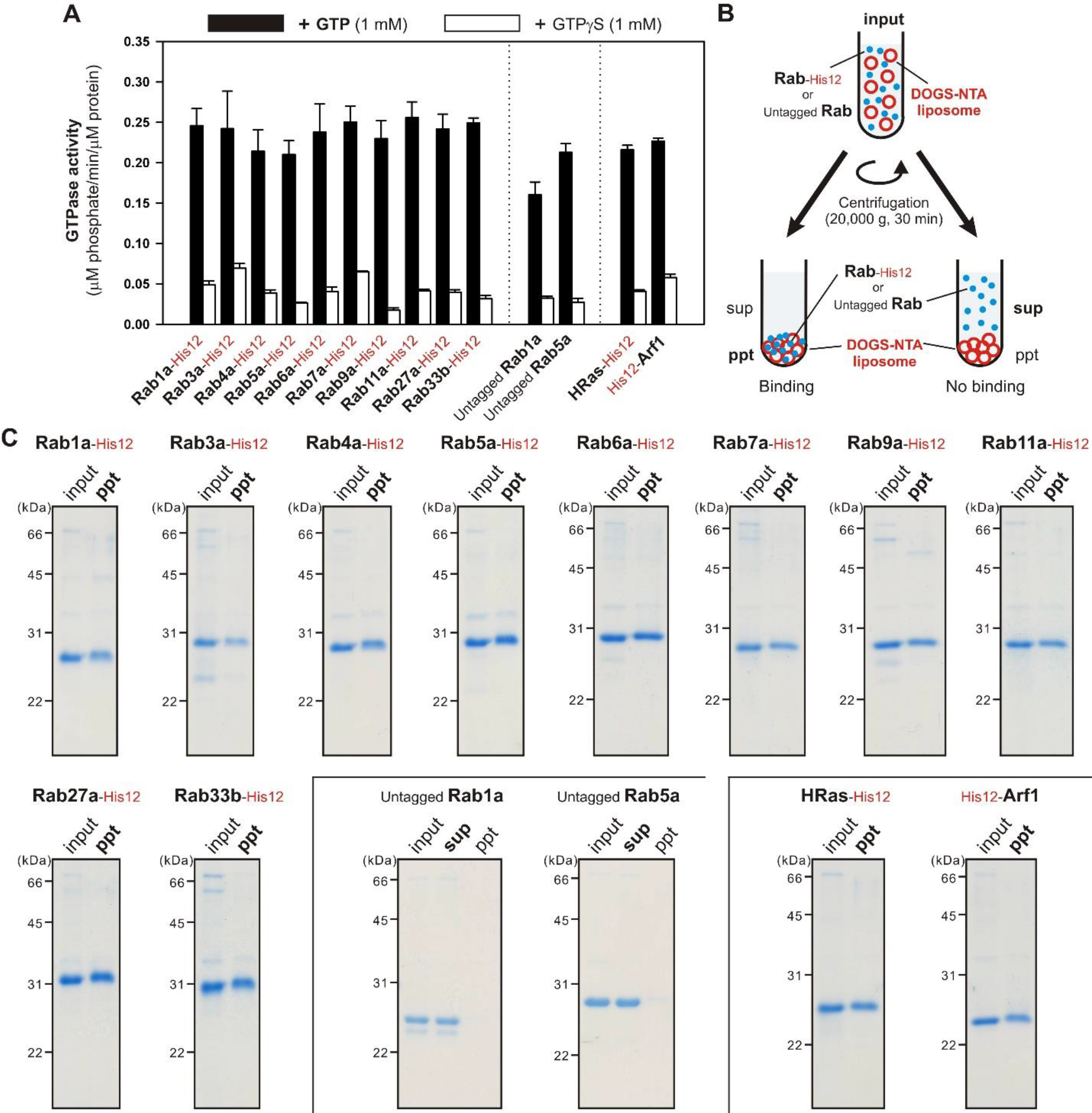
Biochemical characterization of purified Rab-family proteins tested in reconstituted membrane tethering assays. (**A**) GTP-hydrolysis activities of purified recombinant proteins of Rab-His12, untagged Rab, HRas-His12, and His12-Arf1 small GTPases. Purified small GTPases (2 μM) were incubated with 1 mM GTP or GTPγS in RB150 containing 6 mM MgCl_2_ and 1 mM DTT (30°C, 1 h), followed by assaying the free phosphate molecules released in the hydrolytic reactions and determining the specific GTPase activities (μM phosphate/min/μM protein). (**B**) Schematic representation of liposome co-sedimentation assays testing the membrane association of human Rab-family proteins. (**C**) Liposome co-sedimentation assays with purified Rab-His12, untagged Rab, HRas-His12, and His12-Arf1 proteins. Purified small GTPases (4 μM) were incubated with DOGS-NTA-bearing liposomes (2 mM lipids; 1000 nm diameter) in RB150 containing 5 mM MgCl_2_ and 1 mM DTT (30°C, 30 min), centrifuged (20,000 × *g*, 30 min, 4°C), and analyzed via SDS-PAGE for precipitates (ppt) and supernatants (sup) obtained after centrifugation.

### Homotypic *trans*-assembly of human Rab-family proteins in Rab-mediated membrane tethering

Several recent *in vitro* studies using chemically defined reconstitution approaches revealed the intrinsic membrane tethering potency of Rab-family small GTPases in the yeast *Saccharomyces cerevisiae* [26] and in humans [8, 27, 28], leading to a new working model in which membrane-anchored Rab-family proteins function as a *bone fide* membrane tether to physically link two distinct lipid bilayers together during eukaryotic membrane trafficking events [8]. Nevertheless, since Rab-family proteins have been generally believed to only support membrane recruitment of the so-called tethering factors, such as coiled-coil tethering proteins and multisubunit tethering complexes, not to directly promote tethering, in the current paradigm [1, 5–7], it is critical to thoroughly consider the physiological significance of Rab-mediated membrane tethering reactions observed in a chemically defined *in vitro* system. To address this issue, we further explored membrane reconstitution studies on human Rab-mediated membrane tethering by developing a new high-throughput 384-well microplate-based assay to monitor the turbidity of liposomes (Figure 3A), and using the high-throughput turbidity assays to comprehensively and quantitatively test the intrinsic tethering activities of the 10 representative human Rabs and other small GTPases (Figure 1) at a wide range of protein concentrations (0.125-16 μM) (Figure 3B-M). In the turbidity assays, we used the synthetic liposomes (400 nm in diameter) roughly mimicking subcellular compartments in mammalian cells and bearing five major lipid species: phosphatidylcholine (PC), phosphatidylethanolamine (PE), phosphatidylinositol (PI), phosphatidylserine (PS), and cholesterol (Figure 3A) [35]. Nevertheless, it should be noted that the lipid composition used lacks minor but potentially important lipid species, such as phosphoinositides, which may be functional during Rab-dependent membrane tethering events [11, 18]. Rab protein densities on liposomal membrane surfaces, which are correlated with Rab protein-to-lipid molar ratios, are also a critical factor for establishing physiologically relevant conditions in reconstituted membrane tethering reactions (Figure 3B-M), as previously studied on SNARE protein densities in reconstituted SNARE-mediated lipid mixing and membrane fusion [36–39]. Using the average copy number of Rab molecules (25 Rab proteins per vesicle) determined by proteomic and lipidomic analyses of synaptic vesicles from rat brain [40], the mean outer diameter of synaptic vesicles (42 nm) [40], the typical thickness of biological membranes (4 nm) [41], and the average surface area of the headgroups of phospholipids (0.65 nm^2^) [41], we can calculate the Rab-to-lipid molar ratio of synaptic vesicles as 1:560 (25 Rabs/vesicle to 14,100 lipids/vesicle); we thereby refer to this value as the physiological Rab-to-lipid molar ratio in this study, as represented in the liposome turbidity assay data (Figure 3B-M). Although we used the protein stoichiometry of synaptic vesicles as a reasonable model for calculating the physiological Rab protein density [40], it is also conceivable that the Rab densities on other subcellular membrane compartments are variable and significantly different from the current estimations. Strikingly, of the 10 representative human Rabs selected, only early endosomal Rab5a retained the intrinsic capacity to directly trigger the tethering of liposomal membranes below the physiological Rab-to-lipid ratio of 1:560 (mol/mol), and did so at a ratio ranging from 1:800 (at 1 μM Rab5a) to 1:6,400 (at 0.125 μM Rab5a) (Figure 3E). Three other human Rabs, exocytotic Rab3a (Figure 3C), Golgi-resident Rab6a (Figure 3F), and late endosomal/lysosomal Rab7a (Figure 3G), also exhibited their inherent high potency to drive efficient membrane tethering by themselves at the Rab-to-lipid ratios of 1:400 (at 2 μM Rabs), which is close to the physiological ratio of 1:560. In addition, when tested at the higher Rab densities with the Rab-to-lipid ratios of 1:200 (at 4 μM Rabs) and 1:100 (at 8 μM Rabs), another five human Rabs, Rab1a (Figure 3B), Rab4a (Figure 3D), Rab9a (Figure 3H), Rab11a (Figure 3I), and Rab33b (Figure 3K), showed significant tethering activities under these physiologically relevant conditions. However, unlike these 9 Rab-family proteins tested (Figure 1A), Rab27a demonstrated little or no tethering activity, even at Rab-to-lipid ratios of up to 1:50 (at 16 μM Rabs) (Figure 3J), the same activity as the non-Rab Ras-superfamily small GTPases HRas (Figure 3L) and Arf1 (Figure 3M), which were used for the negative controls. Considering that Rab27a exhibits an unconventional Rab isoform that has a ten-residue loop region inserted between two β-strands (Ala55-Arg64) (Figure S1) [32, 42], these experimental data from the current comprehensive and quantitative turbidity assays faithfully reflect that the membrane tethering capacities observed represent a specific molecular function conserved throughout the majority of Rab-family small GTPases (except for Rab27a), and yet their tethering potency appears to be variable for each of the Rab isoforms (Figure 3B-M). It should be also noted that, assuming that all of the Rab-family proteins under investigation are stably bound to liposomal membranes and are typically spherical 25-kDa proteins with an average radius of 2.0 nm [8, 31, 43], the membrane-anchored Rab proteins occupy only 0.59% (at 0.125 μM Rabs) to 38% (at 8 μM Rabs) of the outer membrane surface areas when tested at physiologically relevant Rab-to-lipid ratios (1:6,400-1:100) in the reconstituted membrane tethering assays (Figure 3B-M). This suggests that the human Rab proteins in the tethering reactions have adequate space to physically associate with lipid molecules and other proteins on the membrane surfaces under the current experimental conditions used (Figure 3A).

**Figure 3.**
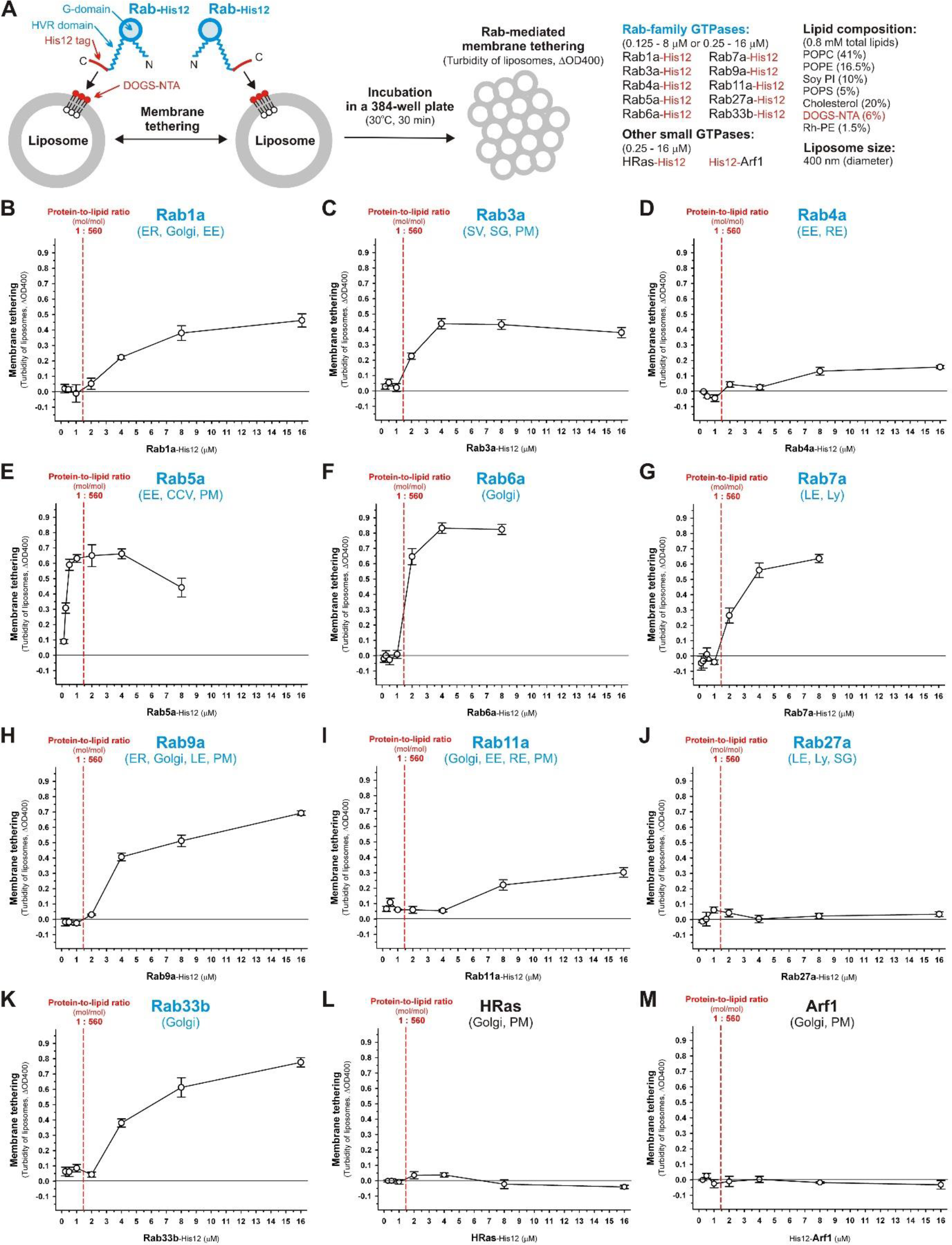
Intrinsic potency of human Rab-family proteins to drive homotypic membrane tethering in a chemically defined reconstitution system. (**A**) Schematic representation of liposome turbidity assays testing homotypic membrane tethering mediated by human Rab-family proteins. (**B-M**) Comprehensive and quantitative analysis of homotypic Rab-mediated membrane tethering by the endpoint liposome turbidity assays for human Rab1a (**B**), Rab3a (**C**), Rab4a (**D**), Rab5a (**E**), Rab6a (**F**), Rab7a (**G**), Rab9a (**H**), Rab11a (**I**), Rab27a (**J**), Rab33b (**K**), HRas (**L**), and Arf1 (**M**). The C-terminal or N-terminal His12-tagged Rab-family proteins, HRas, and Arf1 (each at 0.125, 0.25, 0.5, 1, 2, 4, 8, or 16 μM) were incubated with DOGS-NTA-bearing liposomes (0.8 mM lipids, 400 nm diameter) in RB150 containing 5 mM MgCl_2_ and 1 mM DTT in a 384-well microplate (30°C, 30 min). The turbidity changes in the Rab-mediated liposome tethering reactions were subsequently assayed by measuring the optical density at 400 nm. All turbidity data indicated as ΔOD400 in **B-M**were corrected by subtracting the values of the control liposome reactions in the absence of any Rab-family or other small GTPases. The ΔOD400 values at 8 μM for Rab27a (**J**), HRas (**L**), and Arf1 (**M**) are significantly different from those for all the other nine Rab-family isoforms (**B-I, K**) (*p*<0.001, calculated using one-way ANOVA). The protein-to-lipid molar ratio of 1:560 (mol/mol), shown as red dashed lines in **B-M**, represents the physiological Rab-to-lipid molar ratio, calculated with the copy number of Rab proteins per isolated synaptic vesicle (25 copies per vesicle) [40].

To further explore the efficiency and specificity of homotypic human Rab-mediated membrane tethering in a chemically-defined reconstituted proteoliposome system, the intrinsic membrane tethering capacities of the same 10 representative Rab-family proteins were thoroughly examined by employing fluorescence microscopic observations of Rab-induced liposome clusters (Figure 4). The obtained fluorescence microscopy images were quantitatively analyzed to determine particle numbers, average particle sizes, and total particle areas (Table 1). In the fluorescence microscopy assays of reconstituted membrane tethering, rhodamine (Rh)-labeled 800-nm liposomes (2 mM lipids final) were incubated with purified human Rab-His12 proteins (each at 8 μM final; 1:250 Rab-to-lipid molar ratios) (Figure 4A) and subsequently applied to a LUNA-FL fluorescence cell counter (Logos Biosystems), which allowed us to acquire the wide-field fluorescence images of the Rab-mediated tethering reactions in a defined volume (length × width × height = 2,500 μm × 2,000 μm × 100 μm) (Figure 4B-N), thereby ensuring an unbiased quantitative particle-size analysis of the liposome clusters formed (Table 1). Consistent with the results obtained from the liposome turbidity assays (Figure 3), all of the Rab isoforms tested except for Rab27a (Rab1a, 3a, 4a, 5a, 6a, 7a, 9a, 11a, and 33b) exhibited their intrinsic membrane tethering activities in the microscopy assays, significantly promoting the formation of liposome clusters (Figure 4C-L, Table 1), whereas no or few liposome clusters were observed when assayed without any small GTPase proteins or with Rab27a, HRas, and Arf1 (Figure 4B, K, M, N, Table 1). Remarkably, the current quantitative particle-size analysis revealed a huge difference in the tethering potency among the Rab isoforms. For example, when comparing the two early endosomal Rab isoforms, Rab4a and Rab5a, there were over 20-fold and 300-fold differences in the average particle size and total area of particles, respectively (Table 1). These quantitative imaging data further establish that there is a wide diversity in the intrinsic tethering potency among the human Rab-family proteins (Figure 4, Table 1), even though all of the Rab isoforms share the conserved Ras-superfamily GTPase domain as a major portion in their amino acid sequences (Figure 1A, S1) and indeed exhibit comparable GTP-hydrolysis activities (Figure 2A). This led us to postulate that a specific sequence motif critical to the control of the tethering functions of human Rabs is encoded in the unconserved flexible region at the C-terminus (termed HVR), the N-terminal unconserved flexible segment, and/or the four Rab subfamily-specific regions (termed RabSF1, RabSF2, RabSF3, and RabSF4) that reside within or adjacent to the Ras-superfamily G-domain (Figure 1A, S1) [32, 42, 44].

**Table 1.** Particle size analysis of liposome clusters induced by Rab-mediated membrane tethering. ^*a*^.

| Protein added | Particle number <sup>b</sup> | Average particle size <sup>b</sup><br>($\mu\text{m}^2$) | Total area of particles <sup>b</sup><br>($\mu\text{m}^2$) |
| --- | --- | --- | --- |
| No Rab | 8.7 ± 0.58 | 40 ± 4.3 | 340 ± 22 |
| Rab1a | 2,700 ± 250 | 50 ± 2.9 | 140,000 ± 21,000 |
| Rab3a | 630 ± 50 | 950 ± 83 | 590,000 ± 6,800 |
| Rab4a | 55 ± 14 | 38 ± 4.3 | 2,100 ± 740 |
| Rab5a | 880 ± 140 | 910 ± 150 | 780,000 ± 14,000 |
| Rab6a | 400 ± 55 | 1,700 ± 260 | 660,000 ± 22,000 |
| Rab7a | 460 ± 120 | 1,600 ± 300 | 700,000 ± 78,000 |
| Rab9a | 3,400 ± 35 | 140 ± 4.4 | 470,000 ± 15,000 |
| Rab11a | 190 ± 91 | 33 ± 2.8 | 6,000 ± 2,300 |
| Rab27a | 20 ± 7.8 | 41 ± 9.8 | 860 ± 460 |
| Rab33b | 3,000 ± 410 | 160 ± 13 | 460,000 ± 34,000 |
| HRas | 8.3 ± 6.7 | 45 ± 15 | 440 ± 470 |
| Arf1 | 7.0 ± 3.6 | 98 ± 16 | 650 ± 230 |

<sup>a</sup> Particle sizes of the Rab-induced liposome clusters observed in fluorescence microscopic images were quantitatively measured using ImageJ2 software.
<sup>b</sup> Means and standard deviations of particle numbers, average particle sizes, and total areas of the Rab-induced liposome clusters were determined from three independent rhodamine fluorescence images.

**Figure 4.**
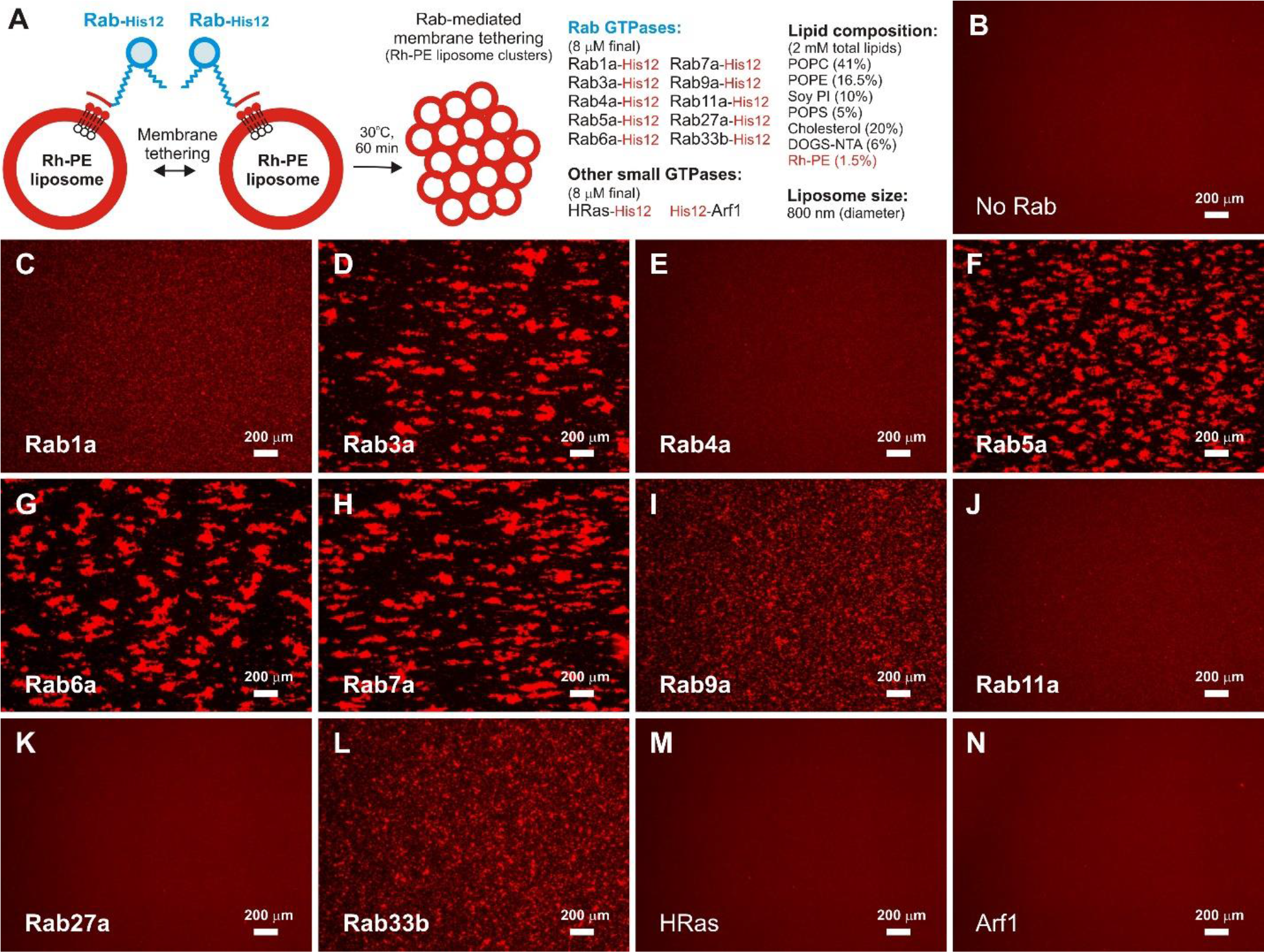
Fluorescence microscopy analysis of liposome clusters induced by homotypic Rab-mediated membrane tethering. (**A**) Schematic representation of fluorescence microscopy analysis of Rab-induced liposome clusters. (**B-N**) Fluorescence images of Rab-induced liposome clusters. Fluorescence-labeled liposomes bearing Rh-PE and DOGS-NTA (2 mM lipids; 800 nm diameter) were incubated in RB150 containing 5 mM MgCl_2_ and 1 mM DTT (30°C, 60 min) in the absence (**B**) and the presence of Rab1a-His12 (**C**), Rab3a-His12 (**D**), Rab4a-His12 (**E**), Rab5a-His12 (**F**), Rab6a-His12 (**G**), Rab7a-His12 (**H**), Rab9a-His12 (**I**), Rab11a-His12 (**J**), Rab27a (**K**), Rab33b-His12 (**L**), HRas-His12 (**M**), and His12-Arf1 (**N**) (each at 8 μM final). After incubation, the Rab-mediated tethering reactions were subjected to fluorescence microscopy and the obtained rhodamine fluorescence images were quantitatively analyzed (Table 1). Scale bars: 200 μm.

Based on our earlier studies on membrane attachment of human Rab-family proteins in reconstituted membrane tethering reactions [27], we revisited the question of whether homotypic Rab-mediated membrane tethering is primarily driven by *trans*-assembly between membrane-associated Rab proteins on two distinct opposing membranes (Figure 5). To address this issue, our analysis employed quantitative approaches with fluorescence cell counter-based imaging assays (Figure 5A-C, F, G) and 384-well microplate-based turbidity assays (Figure 5D, E). Early endosomal Rab5a-His12, which was used as a model for typical human Rabs, was first tested using the microscopic imaging assays with two types of fluorescence-labeled liposomes bearing either Rh-PE or FL-PE (fluorescein-PE) and a DOGS-NTA lipid for anchoring Rab5a-His12 on the membrane surfaces (Figure 5A-C). As expected, Rab5a-anchored Rh-PE liposomes, which were visualized in red (middle panel, Figure 5A), and FL-PE liposomes, which were visualized in green (right panel, Figure 5A), were both able to form massive liposome clusters, yielding total particle areas of 640,000 ± 23,900 μm^2^ and 563,000 ± 53,300 μm^2^, respectively; further both liposomes almost perfectly overlapped with the liposome aggregates observed in the bright-field image (left panel, Figure 5A). However, when a DOGS-NTA lipid was omitted from the FL-PE liposomes, the added Rab5a-His12 did not have the potency to cause clustering of the FL-bearing liposomes (right panel, Figure 5B); in contrast, the Rh-PE/DOGS-NTA liposomes in the same tethering reaction still enabled Rab5a-His12 to trigger the formation of large liposome clusters with an average particle size of 669 ± 81.3 μm^2^ and a total particle area of 496,000 ± 31,100 μm^2^ (middle and left panels, Figure 5B). As expected, Rab5a-His12 completely lost its ability to mediate liposome clustering in the absence of a DOGS-NTA lipid in both the Rh-PE liposomes and the FL-PE liposomes (Figure 5C). These results from the imaging assays clearly demonstrated that homotypic human Rab-mediated membrane tethering requires that Rab proteins are attached onto both of the two distinct membrane surfaces destined to be tethered, thereby establishing the requirement of *trans*-assembly between membrane-anchored Rab proteins for homotypic Rab-mediated tethering. Next, to determine whether or not *trans*-assembly of Rab5a proteins in homotypic tethering reactions can be simply mediated by a protein-protein interaction, we employed liposome turbidity assays for the Rab5a-His12-anchored liposomes in the presence of excess amounts of soluble Rab5a lacking a C-terminal His12 tag (untagged Rab5a), which was used as a potential inhibitor to competitively block the *trans*-interactions between membrane-bound Rab5a-His12 proteins (Figure 5D, E). Indeed, homotypic Rab5a-mediated membrane tethering was not significantly inhibited by the addition of untagged Rab5a, even at 64-fold molar excess (at 8 μM untagged Rab5a) over Rab5a-His12 (at 0.125 μM), giving the mean ΔOD400 values of 0.353 ± 0.052 without untagged Rab5a and 0.305 ± 0.023 with 8 μM untagged Rab5a (Figure 5E). Microscopic imaging assays for Rab5a-mediated tethering were also employed in the absence (Figure 5F) and presence (Figure 5G) of untagged Rab5a. Consistent with the results from the turbidity assays (Figure 5E), Rab5a-His12-anchored liposomes retained their high potency to assemble into large clusters in the presence of a 10- to 40-fold molar excess of untagged Rab5a (20 μM final) over Rab5a-His12 (0.5-2 μM final) (Figure 5F, G). Nevertheless, some inhibition by exogenous untagged Rab5a proteins on Rab-mediated liposome clustering was observed when assayed with a 100-fold molar excess of untagged Rab5a (at 20 μM) over Rab5a-His12 (at 0.2 μM) (left panels, Figure 5F, G). Taken together, these reconstituted assays of homotypic Rab-mediated membrane tethering (Figure 5A-C, E-G) provide experimental evidence establishing that membrane-anchored Rab proteins directly and specifically mediate homotypic membrane tethering by exclusively recognizing the membrane-bound forms of Rabs, not the membrane-unbound soluble forms, thereby conferring selective *trans*-Rab-Rab assembly on membranes (Figure 5D).

**Figure 5.**
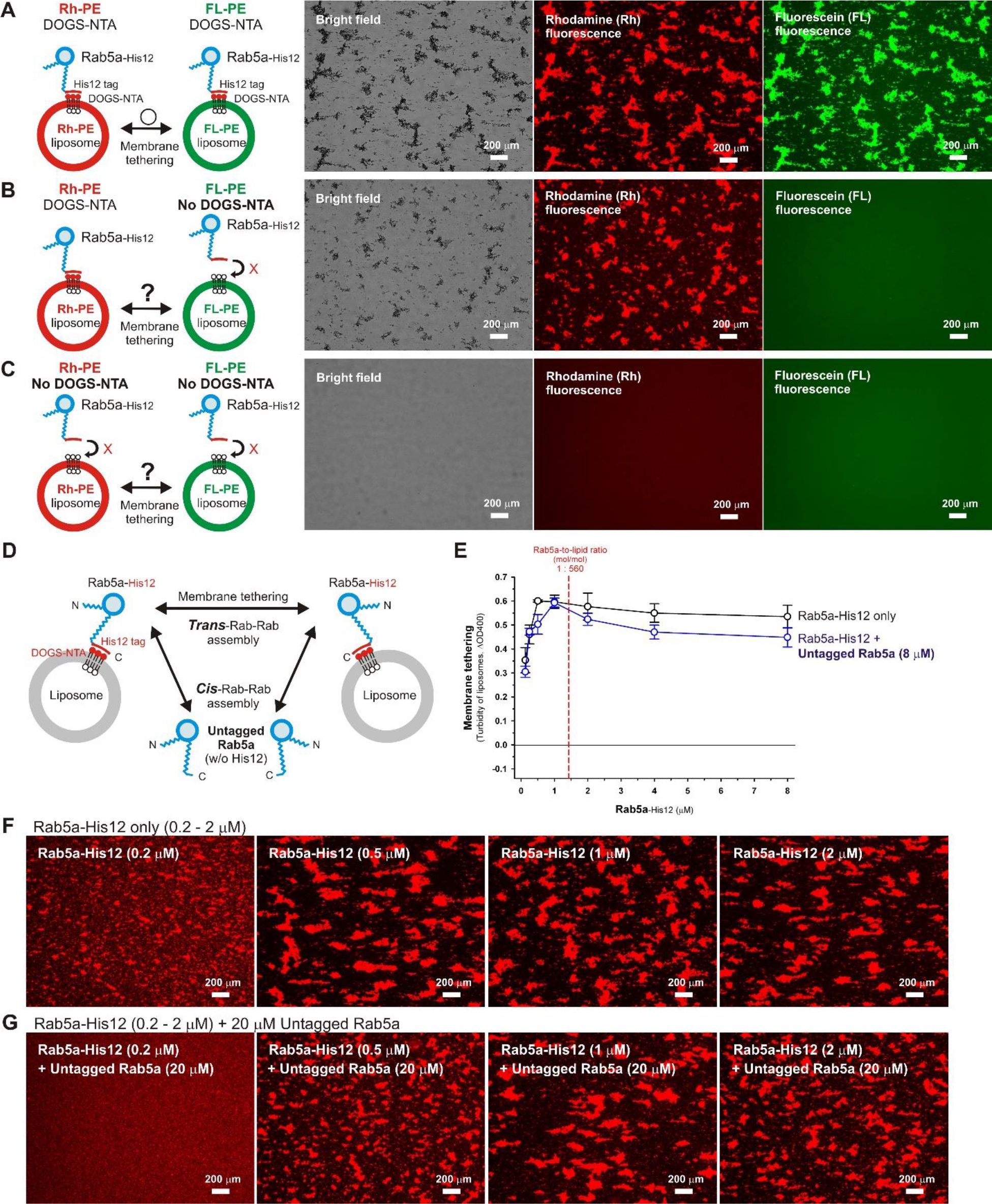
Homotypic Rab-mediated membrane tethering requires *trans*-assembly of Rab proteins on two distinct opposing membranes. (**A-C**) Fluorescence microscopy analysis of homotypic Rab-mediated membrane tethering driven by *trans*-assembly of human Rab5a proteins. Rab5a-His12 (2 μM final) were incubated with two types of fluorescence-labeled liposomes, Rh-PE-bearing liposomes and FL-PE-bearing liposomes (1 mM lipids for each; 800 nm diameter), in RB150 containing 5 mM MgCl_2_ and 1 mM DTT (30°C, 60 min), followed by fluorescence microscopy to obtain bright field images, rhodamine fluorescence images, and fluorescein fluorescence images of the tethering reactions. DOGS-NTA lipids were present in both the Rh-PE and FL-PE liposomes (**A**), only in Rh-PE liposomes (**B**), or not present in either of the liposomes (**C**). Scale bars: 200 μm. (**D**) Schematic representation of a *trans*-assembly of Rab5a proteins on two distinct membranes and a *cis*-assembly of Rab5a proteins on one membrane. (**E**) Liposome turbidity assays testing homotypic Rab5a-mediated membrane tethering in the presence of an excess amount of untagged Rab5a in solution. Rab5a-His12 (0.125, 0.25, 0.5, 1, 2, 4, or 8 μM final) and untagged Rab5a (8 μM final) were preincubated in RB150 containing 5 mM MgCl_2_ and 1 mM DTT (30°C, 10 min), mixed with DOGS-NTA-bearing liposomes (0.8 mM lipids in final; 400 nm diameter), further incubated (30°C, 30 min), and assayed for turbidity changes as in Figure 3 (blue open circles). The control reactions without untagged Rab5a (Rab5a-His12 only) were also prepared and assayed as above (black open circles). (**F**, **G**) Fluorescence microscopic observations of Rab5a-induced liposome clusters in the presence of excess untagged Rab5a. Rab5a-His12 (0.2, 0.5, 1, or 2 μM final) and untagged Rab5a (20 μM final) were preincubated in RB150 containing 5 mM MgCl_2_ and 1 mM DTT (30°C, 10 min), mixed with fluorescence-labeled liposomes bearing Rh-PE and DOGS-NTA (2 mM lipids in final; 800 nm diameter), further incubated (30°C, 30 min), and subjected to fluorescence microscopy as in Figure 4. In addition to the tethering reactions with excess untagged Rab5a (**G**), the control reactions without untagged Rab5a (Rab5a-His12 only) were also prepared and tested as above (**F**). Scale bars: 200 μm.

### Heterotypic *trans*-assembly of human Rab isoforms can specifically trigger Rab-mediated membrane tethering in a chemically defined reconstitution system

The present comprehensive and quantitative analyses of “homotypic” Rab-mediated membrane tethering in a chemically defined reconstitution system (Figure 3–5, Table 1) establish that, in a physiological context, homotypic membrane tethering can be directly driven by *trans*-assembly of human Rab-family isoforms. Specifically, Rab3a, Rab5a, Rab6a, and Rab7a exhibited significant or high intrinsic tethering activities at physiological Rab-to-lipid molar ratios (Figure 3C, E-G), whereas other Rab isoforms were rather inefficient (Rab1a, Rab4a, Rab9a, Rab11a, and Rab33b) or almost incompetent (Rab27a) in initiating homotypic membrane tethering (Figure 3B, D, H-K). Intriguingly, the two Rab isoforms that demonstrated high tethering potency, Rab5a and Rab7a, have been reported to participate in homotypic membrane tethering and fusion events in intracellular endocytic trafficking pathways, such as homotypic fusion of Rab5-positive early endosomes and fusion between Rab7-positive late endosomal and lysosomal compartments [18, 19]. These findings have led to speculation that, in addition to homotypic tethering, heterotypic tethering processes in exocytotic and endocytic pathways can be mediated by heterotypic combinations of Rab-family proteins. Thus, even the Rab isoforms exhibiting low or modest tethering activities in homotypic tethering processes could be more competent for driving efficient membrane tethering in a heterotypic fashion. To test this hypothesis, we next employed 384-well microplate-based liposome turbidity assays in the presence of heterotypic combinations of Rab-family proteins, using the eight Rab pairs as a model (Figure 6A). The assays included Rab1a (1 μM final, Rab-to-lipid ratio of 1:800) and the other eight Rab isoforms paired (each at 0.25-8 μM final, Rab-to-lipid ratios of 1:3,200-1:100), which include Rab3a (Figure 6B), Rab4a (Figure 6C), Rab6a (Figure 6D), Rab7a (Figure 6E), Rab9a (Figure 6F), Rab11a (Figure 6G), Rab27a (Figure 6H), and Rab33b (Figure 6I). Strikingly, of the eight heterotypic Rab combinations containing Rab1a, which is known to be primarily located on the endoplasmic reticulum (ER) and Golgi apparatus (Golgi) compartments (Figure 1), we found that the ER/Golgi-localized Rab1a isoform can selectively cooperate with two of the Golgi-resident isoforms, Rab6a (Figure 6D) and Rab9a (Figure 6F), to synergistically trigger efficient membrane tethering reactions. In the presence of Rab1a (at 1 μM), Rab6a and Rab9a (each at 1 μM) exhibited their high intrinsic capacities to drive membrane tethering even below physiological Rab-to-lipid molar ratios (blue open circles, Figure 6D, F). However, these two Golgi-related Rab isoforms had no potency to cause membrane tethering at the same concentrations when Rab1a was not present in the reactions (black open circles, Figure 6D, F). In contrast, for the other heterotypic Rab combinations tested, Rab1a was unable to significantly stimulate membrane tethering reactions (Figure 6B, C, E, G, H, I), and appeared to rather inhibit the tethering reactions with exocytotic Rab3a and endosomal Rab4a isoforms (blue open circles, Figure 6B, C). These results from the comprehensive turbidity assays provide support for the direct and specific promotion of Rab-mediated membrane tethering by heterotypic combinations of human Rab-family isoforms in a chemically defined reconstitution system.

**Figure 6.**
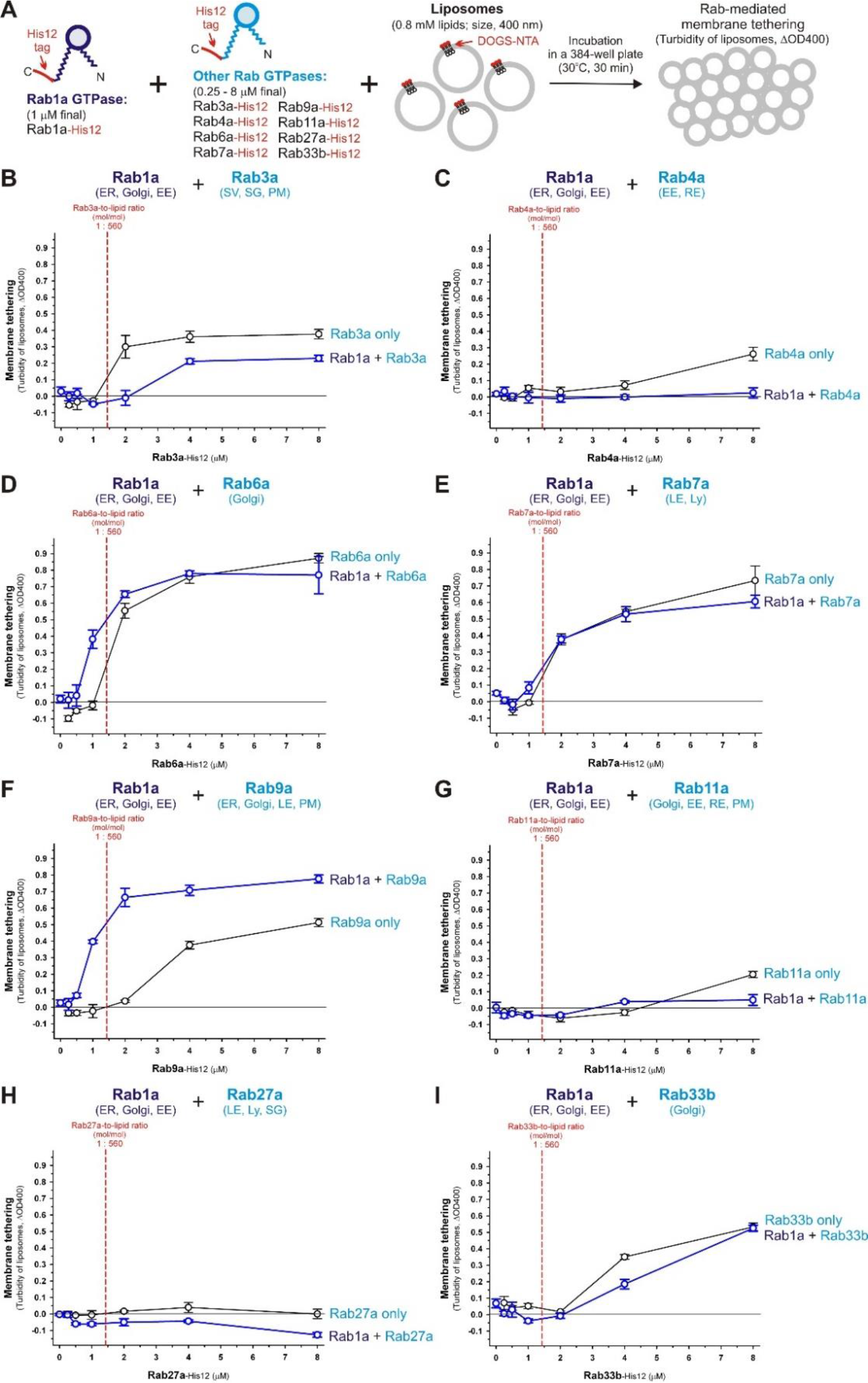
Heterotypic combinations of human Rab-family proteins can synergistically trigger efficient membrane tethering in a chemically defined reconstitution system. (**A**) Schematic representation of liposome turbidity assays with heterotypic combinations of human Rab-family proteins in (**B-I**). (**B-I**) Liposome turbidity assays testing membrane tethering mediated by heterotypic combinations of Rab1a with the other human Rab isoforms. Rab1a-His12 (1 μM final) was preincubated in RB150 containing 5 mM MgCl_2_ and 1 mM DTT (30°C, 10 min) with the other Rab isoforms (each at 0.25, 0.5, 1, 2, 4, or 8 μM final), which include Rab3a-His12 (**B**), Rab4a-His12 (**C**), Rab6a-His12 (**D**), Rab7a-His12 (**E**), Rab9a-His12 (**F**), Rab11a-His12 (**G**), Rab27a-His12 (**H**), and Rab33b-His12 (**I**). After preincubation, the reactions containing two different Rab isoforms were mixed with DOGS-NTA-bearing liposomes (0.8 mM lipids in final; 400 nm diameter), incubated (30°C, 30 min), and then assayed for turbidity changes as in Figure 3 (blue open circles). The control reactions in the absence of Rab1a-His12 were also incubated with liposomes and assayed for turbidity changes as above (black open circles). The ΔOD400 values at 1 μM for Rab6a (**D**) and Rab9a (**F**), in the presence of Rab1a (blue open circles), are significantly different from those for the same Rab isoforms (**D, F**) but in the absence of Rab1a (black open circles) (*p*<0.001, calculated using one-way ANOVA).

To further strengthen the experimental evidence for “heterotypic” Rab-mediated membrane tethering in a chemically defined reconstitution system, the intrinsic tethering potency of the combination of human Rab1a and Rab9a was thoroughly and quantitatively evaluated by employing kinetic assays for liposome turbidity changes (Figure 7A-D) and fluorescence cell counter-based imaging assays for liposome clustering (Figure 7E-J). Consistent with the preceding results in the endpoint turbidity assays with Rab1a and Rab9a (open blue circles, Figure 6F), these two distinct Rab isoforms synergistically acted upon the reconstituted tethering reactions in kinetic turbidity assays, thereby greatly accelerating the rate of Rab-mediated membrane tethering (Figure 7A-D). In the presence of 2 μM Rab1a, which exhibited no tethering activity by itself (closed black circles, Figure 7B), the initial velocity of membrane tethering at 4 μM Rab9a was over 10-fold higher than that of the Rab9a-only control reaction (open red circles, Figure 7B, C; Figure 7D). Moreover, when Rab1a was present, Rab9a still retained its high tethering potency even at 2 μM (open black circles, Figure 7B; the initial velocity of 0.12 ± 0.0039 ΔOD400/min, Figure 7D), whereas no tethering activity was detected with Rab9a alone (open black circles, Figure 7C). Furthermore, these results in kinetic turbidity assays (Figure 7B-D) were fully supported by alternative data from fluorescence imaging assays with the heterotypic Rab1a/Rab9a combination (Figure 7E-J). The presence of Rab1a (at 2 μM) drastically induced the efficient formation of Rab9a-dependent liposome clusters (Figure 7F, G). Particularly, when tested at a concentration of 4 μM Rab9a, which is close to the physiological Rab-to-lipid molar ratio of 1:560 (mol/mol), the tethering reaction containing both Rab1a and Rab9a exhibited an approximately 10-fold higher particle number (Figure 7H), 20-fold higher average particle size (Figure 7I), and 230-fold higher total particle area (Figure 7J) when compared to those values of the Rab9a-only reaction (Figure 7H-J). Thus, the current quantitative analyses of the Rab1a/Rab9a-mediated membrane tethering reactions demonstrated that Rab1a and Rab9a function interdependently to initiate rapid and efficient membrane tethering, probably through their heterotypic Rab-Rab assembly on the two distinct membrane surfaces destined to be tethered.

**Figure 7.**
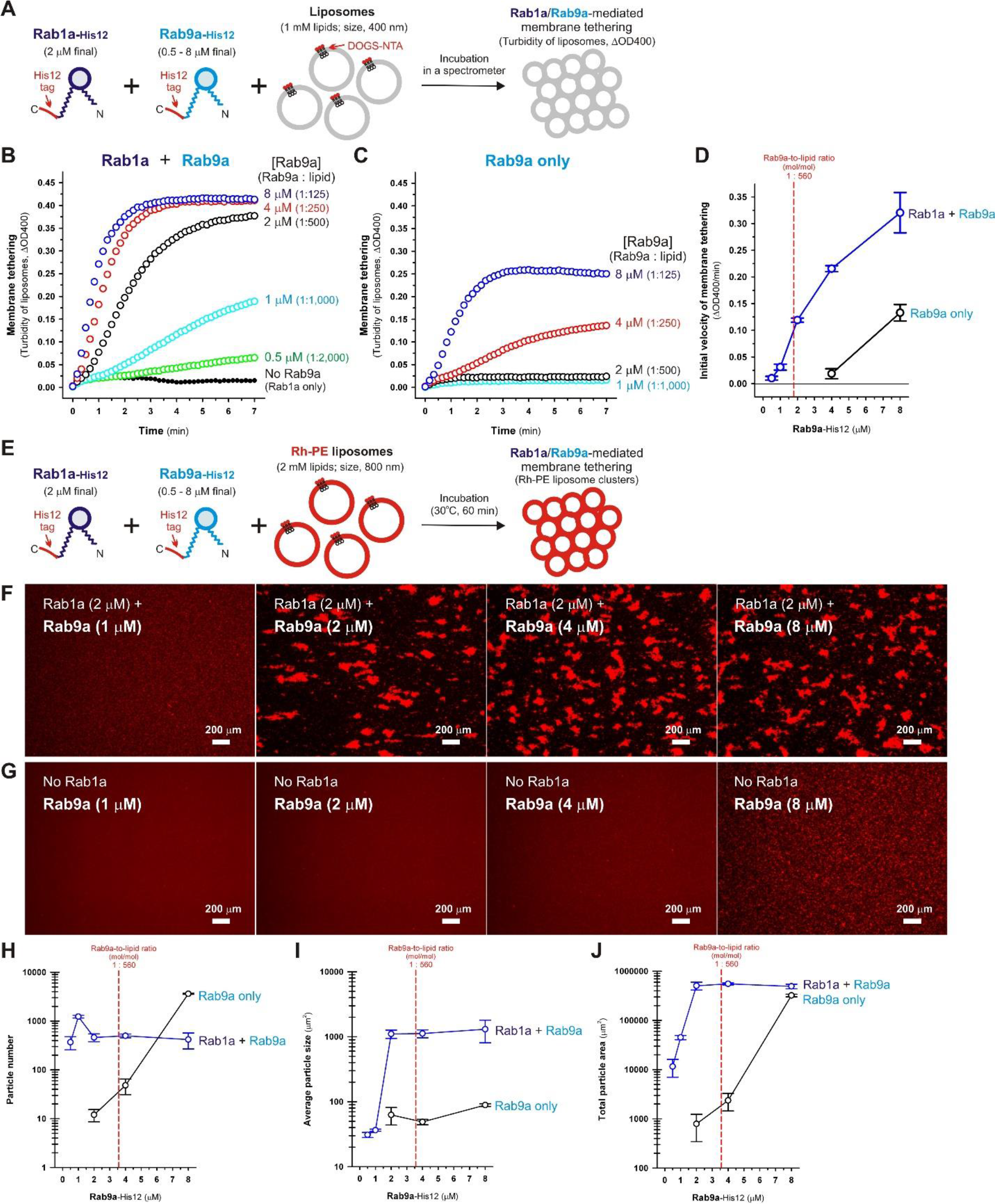
Heterotypic Rab-mediated membrane tethering driven by synergistic actions of human Rab1a and Rab9a proteins. (**A**) Schematic representation of the kinetic liposome turbidity assays with Rab1a-His12, Rab9a-His12, and DOGS-NTA-bearing liposomes in (**B-D**). (**B-D**) Kinetic liposome turbidity assays testing heterotypic membrane tethering mediated by Rab1a and Rab9a. DOGS-NTA-bearing liposomes (1 mM lipids in final; 400 nm diameter) were mixed with Rab9a-His12 (0.5, 1, 2, 4, or 8 μM final) in the presence (**B**) or absence (**C**) of Rab1a-His12 (2 μM final) and immediately assayed for turbidity changes by monitoring the optical density at 400 nm (ΔOD400) for 7 min at 10-sec intervals. The turbidity data obtained were further quantitatively analyzed by curve fitting to determine the initial velocities (ΔOD400/min) of the tethering reactions (**D**). (**E**) Schematic representation of fluorescence microscopic observations of liposome clusters induced by heterotypic membrane tethering mediated by Rab1a and Rab9a. (**F-J**) Fluorescence microscopy of heterotypic Rab1a/Rab9a-mediated membrane tethering reactions. Fluorescence-labeled liposomes bearing Rh-PE and DOGS-NTA (2 mM lipids in final; 800 nm diameter) were mixed with Rab9a-His12 (0.5, 1, 2, 4, or 8 μM final) in the presence (**F**) or absence (**G**) of Rab1a-His12 (2 μM final), incubated (30°C, 60 min), and subjected to fluorescence microscopic observations as in Figure 4. Scale bars: 200 μm. The rhodamine fluorescence images obtained were quantitatively analyzed using ImageJ2 software, yielding means and standard deviations of the particle numbers (**H**), average particle sizes (**I**), and total particle areas (**J**), which were determined from three independent images of the tethering reactions with Rab1a/Rab9a (blue open circles) and Rab9a only (black open circles).

Next, we sought to determine whether heterotypic “*trans*”-assembly of human Rab1a and Rab9a proteins between two opposing membranes was indeed required and also sufficient for specifically driving reconstituted membrane tethering. This was investigated by exploring the intrinsic tethering capacities of pre-incubated Rab1a-anchored liposomes (Rab1a-LPs) and Rab9a-anchored liposomes (Rab9a-LPs) using kinetic liposome turbidity assays (Figure 8A-D) and fluorescence cell counter-based imaging assays (Figure 8E-I). After separately pre-incubating two distinct sets of liposomes anchoring only a single Rab isoform, Rab1a-LPs and Rab9a-LPs (Figure 8A), these Rab-anchored liposomes were mixed and immediately assayed for turbidity changes in the liposome suspensions, thereby exhibiting their high potency to directly trigger heterotypic Rab-mediated membrane tethering (open blue circles, Figure 8B). Efficient and rapid tethering between Rab1a-LPs and Rab9a-LPs was completely abolished by omitting either of the two Rab isoforms from the membrane surfaces (black and green lines, Figure 8B), but was not inhibited at all by the addition of a large molar excess of soluble Rab1a lacking a C-terminal His12 tag (untagged Rab1a) (open red circles, Figure 8B), which had been tested as a potential inhibitor to competitively block the physical interactions between the membrane-bound forms of Rab1a and Rab9a in *trans* (open red circles, Figure 8B). Thus, these data indicate that heterotypic *trans*-assembly between membrane-anchored Rab1a and Rab9a proteins directly and specifically drive membrane tethering in a chemically defined reconstitution system. The specificity of heterotypic *trans*-assembly of human Rab-family isoforms in reconstituted membrane tethering was further examined by testing the pairs of Rab1a not only with Rab9a, but also with Rab4a, Rab11a, and Rab33b (Figure 8C-I). In contrast to efficient and rapid membrane tethering induced by the Rab1a-LPs and Rab9a-LPs pair (open blue circles, Figure 8D), Rab1a-LPs lost its potency to initiate membrane tethering with either Rab4a-LPs or Rab11a-LPs (black and green lines, Figure 8D), consistent with the prior results of the endpoint liposome turbidity assays (Figure 6C, G). This clearly indicates that, among Rab-family isoforms, Rab1a can recognize and associate in *trans* with Rab9a selectively, conferring the specificity of heterotypic Rab-mediated membrane tethering. However, in addition to Rab9a-LPs, Rab33b-LPs also unexpectedly triggered rapid membrane tethering with Rab1a-LPs (open red circles, Figure 8D), even though Rab33b-dependent liposome tethering was not promoted by the addition of Rab1a in the endpoint turbidity assays (Figure 6I). Fluorescence imaging assays with preincubated Rab-anchored liposomes further confirmed the specific *trans*-assembly of these two heterotypic Rab pairs, Rab1a/Rab9a and Rab1a/Rab33b, in reconstituted membrane tethering (Figure 8E-I). Rh-labeled Rab1a-LPs were able to selectively form large liposome clusters with FL-labeled Rab9a-LPs (Figure 8F) and Rab33b-LPs (Figure 8I), whereas Rab1a-LPs had no potency to induce detectable particles of fluorescence-labeled liposomes when tested with Rab4a-LPs (Figure 8G) and Rab11a-LPs (Figure 8H). It is intriguing and noteworthy that Rab33b exhibited its ability to mediate heterotypic tethering with Rab1a exclusively in the reconstituted assays using the pre-incubated liposomes bearing a single Rab isoform (Figure 8D, I). Because both Rab33b and Rab1a are required to be present simultaneously on the same membrane surface in the endpoint turbidity assays (Figure 6A, I), it is conceivable that Rab33b and Rab1a may stably associate with each other in *cis* on one membrane, rather than in *trans* between two distinct membranes, preventing them from initiating heterotypic Rab-mediated membrane tethering. Taken together, these data obtained using our chemically defined reconstitution systems establish that heterotypic *trans*-assembly of human Rab-family proteins can function as minimal machinery to directly and specifically drive membrane tethering.

**Figure 8.**
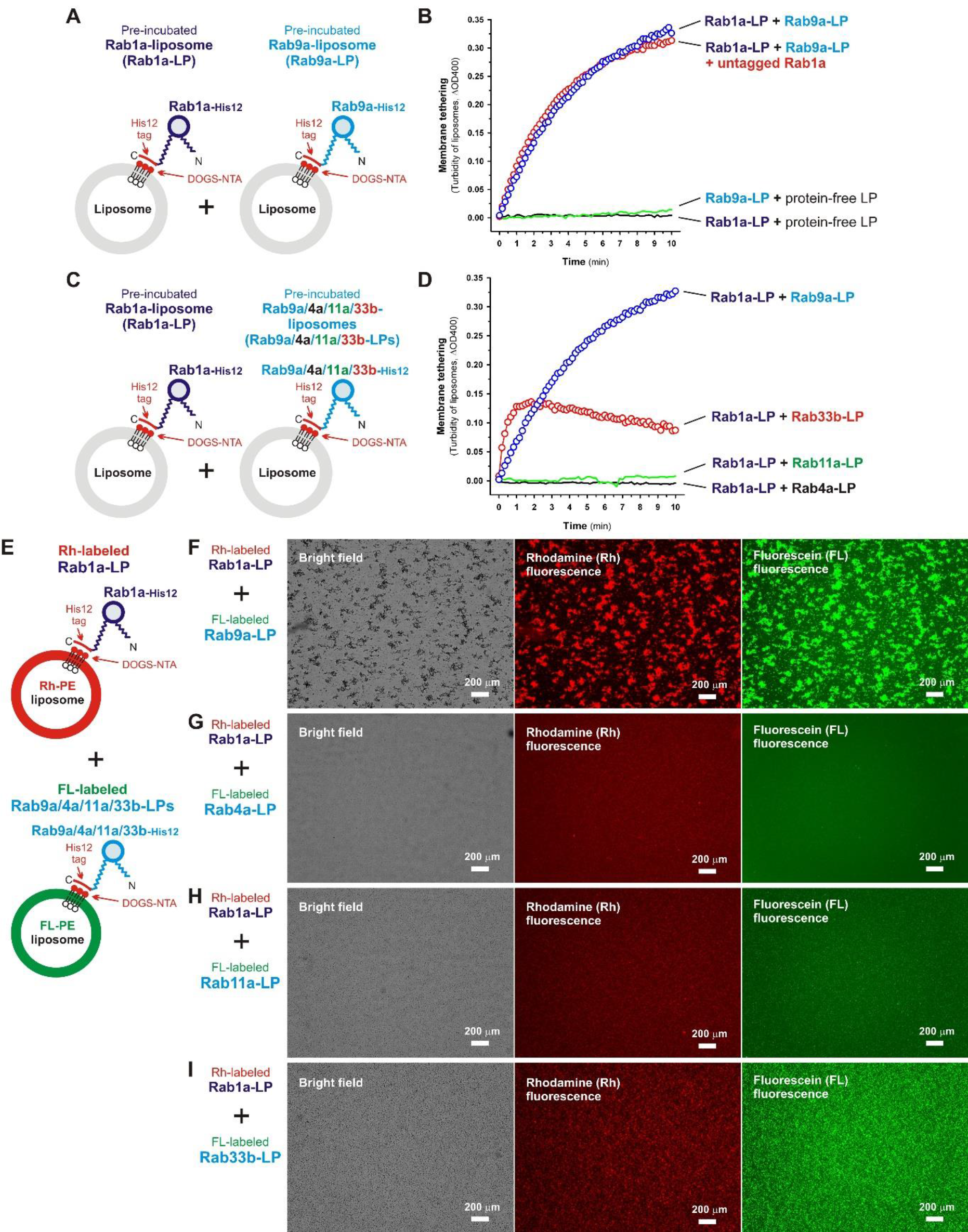
Heterotypic *trans*-assembly of Rab1a and Rab9a on two opposing lipid bilayers specifically drive membrane tethering. (**A**, **B**) Liposome turbidity assays with the pre-incubated Rab1a-anchored liposomes (Rab1a-LPs) and Rab9a-anchored liposomes (Rab9a-LPs), as shown in the illustration (**A**). For the kinetic liposome turbidity assays (**B**), Rab1a-His12 (2 μM final) and Rab9a-His12 (2 μM final) were separately preincubated (30°C, 10 min) with DOGS-NTA-bearing liposomes (1 mM lipids in final; 400 nm diameter) in RB150 containing 5 mM MgCl_2_ and 1 mM DTT to prepare Rab1a-LPs and Rab9a-LPs before starting the turbidity assays (**A**). The pre-incubated Rab1a-LPs and Rab9a-LPs were then mixed and immediately assayed for turbidity changes by monitoring the optical density at 400 nm (ΔOD400) for 10 min at 10-sec intervals (blue open circles). For the controls, the turbidity assays were tested with the reactions in the presence of an excess amount (8 μM final) of untagged Rab1a (red open circles) and the absence of Rab1a-His12 (green line) or Rab9a-His12 (black line). (**C**, **D**) Liposome turbidity assays with pre-incubated Rab-anchored liposomes bearing Rab1a-His12 (Rab1a-LPs), Rab9a-His12 (Rab9a-LPs), Rab4s-His12 (Rab4a-LPs), Rab11a-His12 (Rab11a-LPs), and Rab33b-His12 (Rab33b-LPs), as shown in the illustration (**C**). After preparing the Rab-anchored liposomes, Rab1a-LPs were mixed with Rab9a-LPs (blue open circles), Rab4a-LPs (black line), Rab11a-LPs (green line), or Rab33b-LPs (red open circles), immediately followed by monitoring the turbidity changes (**D**), as described above in (**B**). (**E-I**) Fluorescence microscopy of pre-incubated Rab-anchored liposomes, Rab1a-LPs, Rab9a-LPs, Rab4a-LPs, Rab11a-LPs, and Rab33b-LPs, as shown in the illustration (**E**). The Rab-anchored liposomes were prepared by preincubating Rab-His12 proteins (each at 4 μM final) with fluorescence-labeled liposomes (2 mM lipids in final; 800 nm diameter), which include Rh-PE-bearing liposomes for Rab1a-His12 and FL-PE-bearing liposomes for the other Rab-His12 proteins (**E**), at 30°C for 10 min. The Rh-labeled Rab1a-LPs were mixed with FL-labeled Rab9a-LPs (**F**), Rab4a-LPs (**G**), Rab11a-LPs (**H**), and Rab33b-LPs (**I**), further incubated (30°C, 60 min), and subjected to fluorescence microscopy to obtain bright field images, rhodamine fluorescence images, and fluorescein fluorescence images of the tethering reactions (**F-I**). Scale bars: 200 μm.

### Conclusions

On the basis of the pioneering work of Merz et al., who first reported that endosomal Rab/Ypt small GTPases in the yeast *S. cerevisiae* directly induced membrane tethering *in vitro* [26], and our prior studies on reconstituted human Rab-mediated membrane tethering [8, 27, 28], the present study comprehensively and quantitatively explored the intrinsic capacities of 10 representative human Rab-family isoforms in exocytic and endocytic pathways (Rab1a, 3a, 4a, 5a, 6a, 7a, 9a, 11a, 27a, and 33b; Figure 1, S1) for the physical tethering of two distinct lipid bilayers through their Rab-Rab protein assembly in a homotypic (Figure 3-5) and heterotypic (Figure 6–8) manner. Experimental evidence from the current chemically defined reconstitution systems clearly establishes that both homotypic and heterotypic *trans*-assembly between membrane-anchored human Rab proteins can directly, and solely, trigger efficient and specific membrane tethering under physiologically relevant experimental conditions (Figure 3–8). Notably, endosomal Rab5a retains its particularly high tethering potency, thereby driving efficient homotypic membrane tethering even at densities significantly below the physiological Rab protein density on membrane surfaces, such as the Rab-to-lipid molar ratios of 1:6,400 (Figure 3E, 5E) and 1:10,000 (Figure 5F). As compared to hyperactive Rab5a, the other Rab-family isoforms appear to be relatively inefficient in initiating homotypic and heterotypic Rab-mediated membrane tethering (Figure 3B-M, 6B-I). However, it is conceivable that specific types of Rab-interacting effector proteins or protein complexes can further promote membrane tethering driven by these less active Rab isoforms through direct facilitation of the membrane-bound states and structures of Rabs; for example, we recently discovered that the C-terminal globular tail domains of class V myosins substantially and specifically enhanced the intrinsic tethering capacity of their cognate Rab11a isoform [28]. Additionally, it should be noted that prior reconstitution studies by other groups have established the intrinsic tethering activities of some specific Rab effectors, such as the HOPS tethering complex and the endosomal EEA1 tether, in the presence and/or absence of their cognate Rab-family proteins [22-25]. Thus, how Rab effectors and Rabs cooperate in the process of membrane tethering has yet to be elucidated. Our current findings also highlight the strict selectivity of *trans*-Rab-Rab assembly in homotypic and heterotypic Rab-mediated membrane tethering. A huge molar excess of exogenous soluble Rab5a lacking a membrane anchor (untagged Rab5a) had little or no potency to competitively inhibit the homotypic Rab5a-mediated tethering reactions (Figure 5D-G), thus establishing the stringent recognition and assembly between membrane-bound Rab5a proteins in *trans*. A comprehensive analysis of heterotypic combinations of human Rab-family isoforms in reconstituted membrane tethering reveals that ER/Golgi-resident Rab1a cooperates exclusively with the three Golgi-related Rab isoforms, Rab9a, Rab6a, and Rab33b, to synergistically and specifically trigger rapid and efficient membrane tethering via heterotypic *trans*-Rab-Rab assembly on two opposing membranes (Figure 6–8). Finally, to fully understand how Rab-family small GTPases confer the efficiency and specificity of homotypic and heterotypic Rab-mediated membrane tethering, further studies will be needed to decipher the structural determinants in a Rab protein molecule for *trans*-Rab-Rab assembly on membrane surfaces.

## Experimental procedures

### Protein expression and purification

The expression vectors for human Rab-family small GTPases (Rab1a [UniProtKB: P62820], Rab3a [UniProtKB: P20336], Rab4a [UniProtKB: P20338], Rab5a [UniProtKB: P20339], Rab6a [UniProtKB: P20340], Rab7a [UniProtKB: P51149], Rab9a [UniProtKB: P51151], Rab11a [UniProtKB: P62491], Rab27a [UniProtKB: P51159], and Rab33b [UniProtKB: Q9H082]) and the non-Rab Ras superfamily small GTPases (HRas and Arf1) were constructed using a pET-41 Ek/LIC vector kit (Novagen), as described previously [27, 28]. The coding sequences of these small GTPase proteins were amplified with polymerase chain reaction (PCR) using KOD-Plus-Neo DNA polymerase (Toyobo) and Human Universal QUICK-Clone cDNA II (Clontech) as template cDNA. The oligonucleotide primers used were designed to amplify the PCR fragments containing the additional DNA sequences encoding a human rhinovirus (HRV) 3C protease site (Leu-Glu-Val-Leu-Phe-Gln-Gly-Pro) and polyhistidine residues (His12), which were inserted upstream of the initial ATG codons and downstream of the codons for the C-terminal amino acid residue, respectively. For Arf1 small GTPase as an exception, the His12 coding sequence was located between the upstream sequence for the HRV 3C protease site and the initial ATG codon in the PCR fragment. All of these amplified fragments were cloned into a pET-41 Ek/LIC vector (Novagen) using the ligation-independent cloning method (Novagen), yielding vectors that expressed the N-terminal GST-His6-tagged forms of recombinant Rab-His12, HRas-His12, and His12-Arf1 proteins (Figure 1A). In addition, for Rab1a and Rab5a, the PCR fragments without the His12 coding sequence were also amplified and cloned into the same pET-41-based vector as described above, to prepare Rab1a and Rab5a proteins lacking the C-terminal His12 tag (untagged Rab1a and Rab5a) (Figure 1A).

Recombinant Rab-His12, untagged Rab, HRas-His12, and His12-Arf1 proteins were expressed as the N-terminal GST-His6-tagged form in *Escherichia coli* BL21(DE3) cells (Novagen) harboring pET-41-based vectors in Lysogeny Broth (LB) medium (1 liter each) containing kanamycin (50 μg/ml). After inducing protein expression by adding IPTG (0.1 mM final, 37°C, 3 hours), cultured *E. coli* cells were harvested by centrifugation, resuspended in RB150 buffer (20 mM Hepes-NaOH, pH 7.4, 150 mM NaCl, 10% glycerol) containing 0.1 mM GTP, 5 mM MgCl_2_, 1 mM dithiothreitol (DTT), 1 mM phenylmethylsulfonyl fluoride (PMSF), and 1.0 μg/ml pepstatin A (40 ml each), then freeze-thawed in liquid nitrogen and a water bath at 30°C, lysed by sonication using a UD-201 ultrasonic disrupter (Tomy Seiko), and ultracentrifuged with a 70 Ti rotor (Beckman Coulter; 50,000 rpm, 75 min, 4°C). The supernatants obtained (around 40 ml each) were mixed with COSMOGEL GST-Accept beads (50% slurry, 4 ml each; Nacalai Tesque) and subsequently incubated with gentle agitation (4°C, 2 hours). The protein-bound GST-Accept beads (approximately 2 ml bed volume for each) were washed with RB150 containing 5 mM MgCl_2_ and 1 mM DTT (8 ml each) four times, resuspended in the same buffer (4 ml each) containing HRV 3C protease (4 units/ml final; Novagen), and incubated without agitation (4°C, 16 hours). During the incubation, the full-length Rab-His12, untagged Rab, HRas-His12, and His12-Arf1 proteins, which have only three extra residues (Gly-Pro-Gly) at the N-terminus (Figure 1A), were specifically cleaved off and eluted from the beads by HRV 3C protease cleavage. After centrifugation of the incubated bead suspensions (15,300 × *g*, 10 min, 4°C), the purified proteins were harvested from the supernatants (around 4 ml each). Protein concentrations of the purified Rab-family, HRas, and Arf1 proteins were determined using Protein Assay CBB Solution (Nacalai Tesque) and BSA as a standard protein. The purities of these purified recombinant proteins were analyzed by sodium dodecyl sulfate polyacrylamide gel electrophoresis (SDS-PAGE) and Coomassie blue staining after boiling in 1.6% SDS (100°C, 5 min) (Figure 1B).

### GTPase activity assay

GTPase activities of the purified small GTPase proteins were assayed by quantitating free phosphate molecules released in hydrolytic reactions containing GTP, using the Malachite Green-based reagent Biomol Green (Enzo Life Sciences) as described previously [27, 28]. Purified Rab-His12, untagged Rab, HRas-His12, and His12-Arf1 proteins (2 μM final) were incubated at 30°C for 1 hour in RB150 containing 6 mM MgCl_2_, 1 mM DTT, and 1 mM GTP or GTPγS (100 μl each). After incubation, the hydrolytic reactions (100 μl each) were mixed with the Biomol Green reagent (100 μl each), followed by further incubation at 30°C for 30 min. The reaction mixtures (200 μl each) were then transferred to a clear 96-well microplate (Falcon no. 351172, Corning) and the absorbance at 620 nm (A620) was measured with a SpectraMax Paradigm plate reader (Molecular Devices) using an ABS-MONO cartridge (Molecular Devices). All of the raw A620 data were corrected by subtracting the absorbance values of the control reactions with GTP or GTPγS (1 mM each) in the absence of any small GTPase proteins. To calculate the concentrations of phosphate molecules released in the hydrolytic reactions from the A620 data obtained, phosphate standards (2.5-40 μM final, Enzo Life Sciences) were also incubated in the same buffer and assayed as above. Means and standard deviations of the specific GTPase activities (μM phosphate/min/μM protein) for the purified small GTPases were determined from three independent experiments.

### Liposome preparation

Non-fluorescent lipids used to prepare a synthetic protein-free liposome, including POPC (1-palmitoyl-2-oleoyl-PC), POPE (1-palmitoyl-2-oleoyl-PE), soy PI, POPS (1-palmitoyl-2-oleoyl-PS), DOGS-NTA, and ovine cholesterol, were all purchased from Avanti Polar Lipids. Fluorescence-labeled lipids Rh-PE and FL-PE were obtained from Molecular Probes. Lipid mixes for liposome preparation contained 41% (mol/mol) POPC, 16.5% POPE, 10% soy PI, 5% POPS, 20% cholesterol, 6% DOGS-NTA, and 1.5% fluorescent Rh-PE or FL-PE in chloroform. For the liposomes without DOGS-NTA, POPC was added to the lipid mixes up to 47% (mol/mol). After evaporating chloroform from the lipid mixes with a stream of nitrogen gas, dried lipid films were completely resuspended by vortexing in RB150 containing 5 mM MgCl_2_ and 1 mM DTT (8 mM total lipids in final), incubated at 37°C for 1 hour while shaking, and freeze-thawed in liquid nitrogen and a water bath at 30°C. The lipid suspensions obtained were then extruded 25 times through polycarbonate filters (pore diameters, 400, 800, or 1000 nm; Avanti Polar Lipids) in a mini-extruder (Avanti Polar Lipids) at 40°C. Protein-free liposome solutions thus prepared (8 mM lipids final) were stored at 4°C and used within a week for all of the experiments in the present studies. Size distributions of the prepared liposomes were measured by dynamic light scattering (DLS) using a DynaPro NanoStar DLS instrument (Wyatt Technology) (Figure S2).

### Liposome co-sedimentation assay

To assay the association of Rab-His12, untagged Rab, HRas-His12, and His12-Arf proteins with liposomal membranes, the purified proteins (4 μM final) were mixed with extruded liposomes bearing DOGS-NTA and Rh-PE (1000 nm diameter; 2 mM total lipids in final) in RB150 containing 5 mM MgCl_2_ and 1 mM DTT (150 μl each), subsequently incubated without agitation at 30°C for 30 min, and centrifuged at 20,000 × *g* for 30 min at 4°C. The input samples before incubation and the pellets and supernatants obtained after centrifugation were boiled in 0.7% SDS (100°C, 5 min) and then analyzed using SDS-PAGE and Coomassie blue staining.

### Liposome turbidity assay

The intrinsic membrane tethering capacities of human Rab-family and non-Rab Ras superfamily small GTPases were assayed by measuring turbidity changes in synthetic liposome suspensions in the presence of purified recombinant Rab-family, HRas, and Arf1 proteins, as described previously [8, 27, 28]. For the endpoint assays of homotypic liposome tethering mediated by the small GTPases, DOGS-NTA/Rh-PE-bearing extruded liposomes (400 nm diameter; 0.8 mM total lipids in final) and purified Rab-His12, HRas-His12, or His12-Arf1 proteins (0.125-16 μM in final), which had been separately preincubated (30°C, 10 min), were mixed in RB150 containing 5 mM MgCl_2_ and 1 mM DTT (total 150 μl for each), immediately transferred into a black 384-well plate with a clear flat bottom (40 μl per well; Corning 3544), and incubated without agitation at 30°C for 30 min. After incubation, turbidity changes in the liposome suspensions were measured with optical density values at 400 nm on a SpectraMax Paradigm plate reader using an ABS-MONO cartridge and the PathCheck option (Molecular Devices) that normalizes the obtained values to a 1-cm path length. For the assays testing heterotypic Rab combinations, Rab1a-His12 (1 μM final) was preincubated (30°C, 10 min) with the other Rab-His12 proteins (0.25-8 μM final; Rab3a-His12, Rab4a-His12, Rab6a-His12, Rab7a-His12, Rab9a-His12, Rab11a-His12, Rab27a-His12, or Rab33b-His12) before mixing with liposome suspensions. These liposome turbidity reactions with heterotypic Rab combinations were then incubated in a 384-well plate and assayed for turbidity changes, as described above for the homotypic tethering reactions. All of the turbidity data (ΔOD400) were corrected by subtracting the values of the control liposome reactions without any protein components. Means and standard deviations of the turbidity data (ΔOD400) were determined from three independent experiments.

Kinetic assays of Rab-mediated membrane tethering were also employed by measuring the turbidity changes in liposome suspensions, as described previously [8, 27, 28]. After preincubation at 30°C for 10 min to equilibrate the temperature of the protein and liposome samples used, turbidity reactions were prepared by mixing DOGS-NTA/Rh-PE-bearing liposomes (400 nm diameter; 1 mM lipids final) and purified Rab-His12 proteins (0-10 μM final) in RB150 containing 5 mM MgCl_2_ and 1 mM DTT (total 150 μl for each), immediately followed by applying the reactions to a 10-mm path-length cell (105.201-QS, Hellma Analytics) and monitoring the optical density changes at 400 nm (ΔOD400) for 10 min with 10-sec intervals in a DU720 spectrophotometer (Beckman Coulter) at room temperature. To quantitatively evaluate the initial velocity of Rab-mediated membrane tethering, the turbidity data obtained from the kinetic assays were subjected to curve fitting using ImageJ2 software (National Institutes of Health) and the logistic function formula, y = a/(1+b*exp(−c*x)), in which y and x are the ΔOD400 value and the time (min), respectively. The initial velocities were defined as the maximum slopes of the fitted curves, which can be calculated as c*a/4 from the formula above. Means and standard deviations of the initial velocities (ΔOD400/min) were determined from three independent experiments. All of the kinetic assay data were obtained from one experiment and were typical of those from more than three independent experiments.

### Fluorescence microscopy

Fluorescence microscopy observations of Rab-induced liposome clusters were performed using a LUNA-FL automated fluorescence cell counter (Logos Biosystems) and LUNA cell counting slides (L12001, Logos Biosystems). Fluorescence-labeled liposomes bearing Rh-PE or FL-PE (800 nm diameter; 2 mM total lipids in final) and purified Rab-His12, HRas-His12, or His12-Arf1 proteins (0-10 μM in final) were preincubated separately at 30°C for 10 min, mixed in Rb150 containing 5 mM MgCl_2_ and 1 mM DTT (total 80 μl for each), and further incubated without agitation at 30°C for 60 min, followed by applying liposome suspensions to the cell counting slides (12 μl per well, 2 wells per slide). Bright field images, rhodamine fluorescence images, and fluorescein fluorescence images of the tethering reactions on the slides were obtained and processed using the LUNA-FL fluorescence cell counter. Particle sizes of the liposome clusters observed in rhodamine and fluorescein fluorescence images were measured using ImageJ2 software after setting the lower intensity threshold level to 150, the upper intensity threshold level to 255, and the minimum particle size to 10 pixel^2^ [8, 27, 28].

## Acknowledgements

We thank Motoki Inoshita, Kana Fujibayashi, and Megumi Shinguu (Institute for Protein Research, Osaka University) for contributions to preparing the expression vectors and purified proteins for human Rab-family GTPases and other small GTPases. We are grateful to Drs. Genji Kurisu and Hideaki Tanaka (Institute for Protein Research, Osaka University) for access to dynamic light scattering measurements. This study was in part supported by the Program to Disseminate Tenure Tracking System from the Ministry of Education, Culture, Sports, Science and Technology, Japan (MEXT) and Grants-in-Aid for Scientific Research from MEXT (to J.M.).

## Author contributions

J.M. designed the research; J.M., K.S., and N.T. performed the experiments; J.M., K.S., and N.T. analyzed the data; J.M. wrote the manuscript.

## The abbreviations used are

SNARE: soluble N-ethylmaleimide-sensitive factor attachment protein receptor
HVR: hypervariable region
PC: phosphatidylcholine
PE: phosphatidylethanolamine
PI: phosphatidylinositol
PS: phosphatidylserine
Rh: rhodamine
HRV: human rhinovirus
FL: fluorescein
PO: 1-palmitoyl-2-oleoyl

---

**Figure S1.**
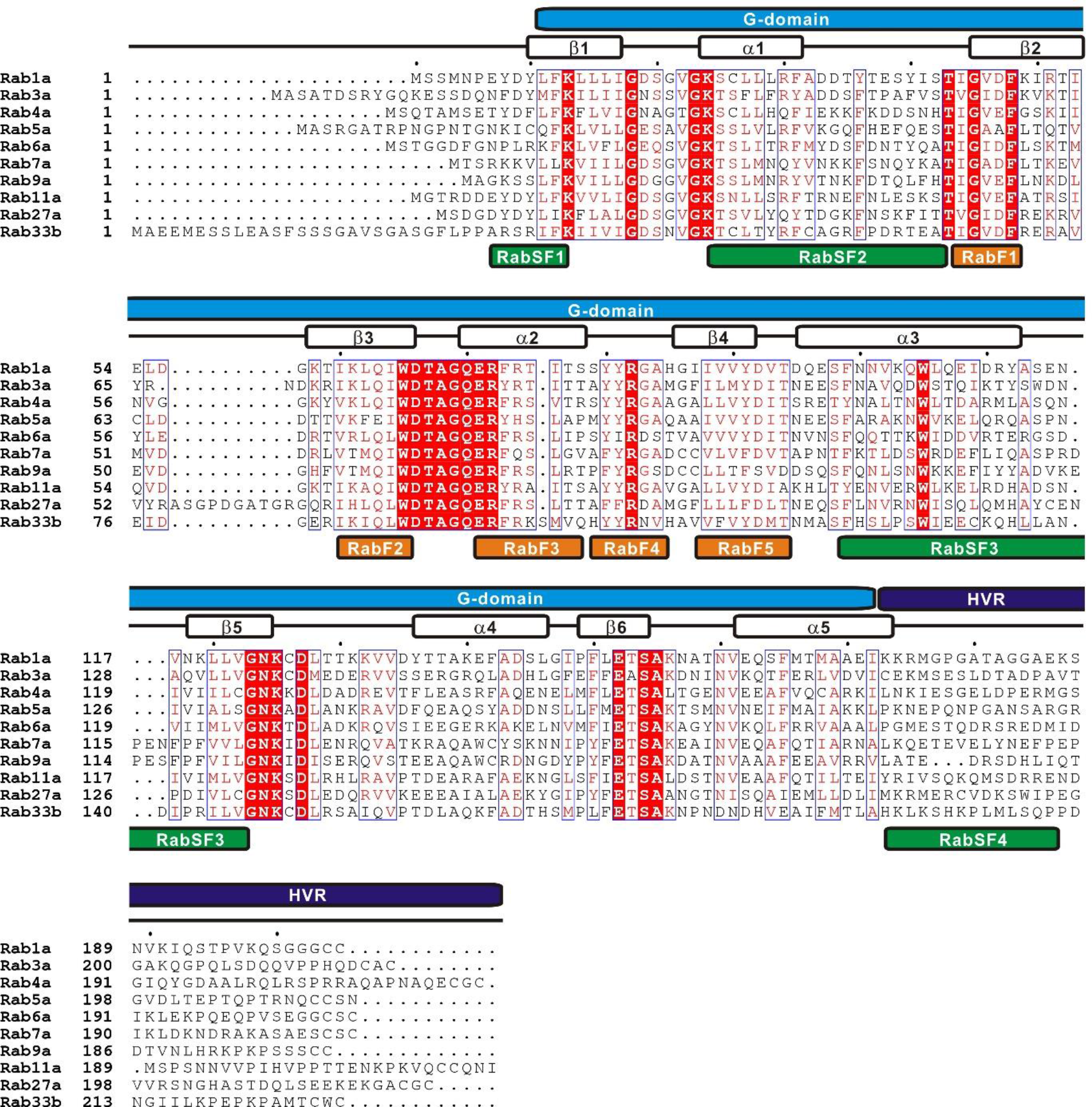
Alignment of amino acid sequences of human Rab-family small GTPases tested in the current study in a chemically defined reconstitution system. Amino acid sequences of the human Rab-family proteins (Rab1a, Rab3a, Rab4a, Rab5a, Rab6a, Rab7a, Rab9a, Rab11a, Rab27a, and Rab33b) were obtained from the UniProtKB knowledgebase (https://www.uniprot.org), aligned using the ClustalW program (https://www.genome.jp/tools-bin/clustalw), and rendered with the ESPript 3.0 program (http://espript.ibcp.fr/ESPript/ESPript/). Identical and similar residues in the sequence alignment are highlighted in a red box and in red characters, respectively. Sequence regions corresponding to the Ras-superfamily GTPase domain (G-domain), hypervariable region (HVR), Rab family specific motifs (RabF1-5), and Rab subfamily specific motifs (RabSF1-4) are indicated on the top or bottom of the sequence alignment [32, 42]. Secondary structures in the Rab3a structure (PDB code, 3RAB), including five α-helices (α1-α5) and six β-strands (β1-β6), are also shown on the top of the sequence alignment.

**Figure S2.**
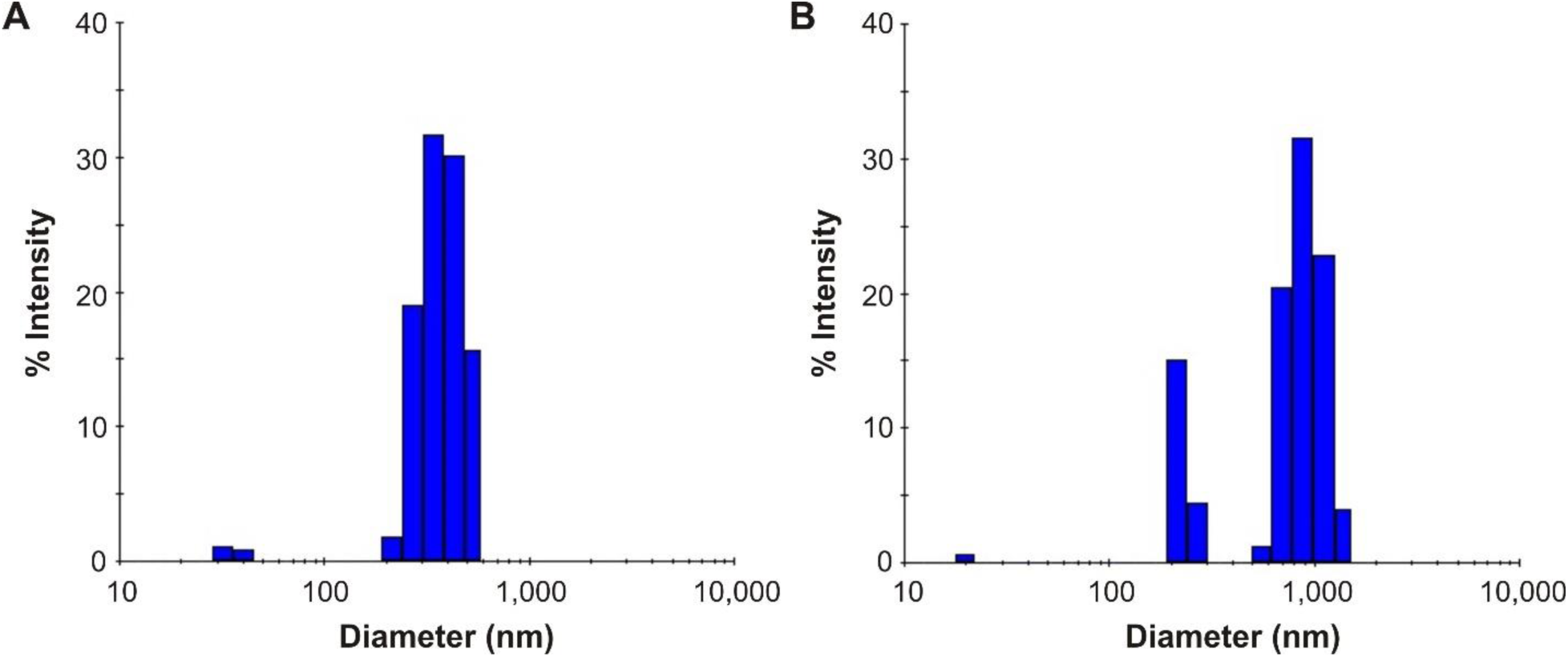
Dynamic light scattering measurements of synthetic liposomes used in the present reconstitution studies. (**A, B**) Histograms of size distributions of typical liposome preparations, Rh-labeled 400-nm liposomes used for turbidity assays (**A**) and Rh-labeled 800-nm liposomes used for fluorescence microscopy experiments (**B**), measured by dynamic light scattering.

